# Evaluation of classifier performance for multiclass phenotype discrimination in untargeted metabolomics

**DOI:** 10.1101/139584

**Authors:** Patrick J. Trainor, Andrew P. DeFilippis, Shesh N. Rai

## Abstract

Statistical classification is a critical component of utilizing metabolomics data for examining the molecular determinants of phenotypes and for furnishing diagnostic and prognostic phenotype predictions in medicine. Despite this, a comprehensive and rigorous evaluation of classification techniques for phenotype discrimination given metabolomics data has not been conducted. We conducted such an evaluation using both simulated and real metabolomics data, comparing Partial Least Squares-Discriminant Analysis (PLS-DA), Sparse PLS-DA, Random Forests, Support Vector Machines, and Neural Network classification techniques for discriminating phenotype. We evaluated the techniques on simulated data generated to mimic global untargeted metabolomics data by incorporating realistic block-wise correlation and partial correlation structures for mimicking the correlations and metabolite clustering generated by biological processes. Over the simulation studies, covariance structures, means, and effect sizes were randomly simulated to provide consistent estimates of classifier performance over a wide range of possible scenarios. The presence of non-normal error distributions and the effect of prior-significance filtering (dimension reduction) were evaluated. In each simulation, classifier parameters (such as the number of hidden nodes in a neural network) were tuned by cross-validation to minimize the probability of detecting spurious results due to poorly tuned classifiers. Classifier performance was then evaluated using real clinical metabolomics datasets of varying sample medium, sample size, and experimental design. We report that in the scenarios without a significant presence of non-normal error distributions over metabolite clusters, Neural Network and PLS-DA classifiers performed poorly relative to Sparse PLS-DA (sPLS-DA), Support Vector Machine (SVM), and Random Forest classifiers. When non-normal error distributions were introduced, the performance of PLS-DA classifiers deteriorated further relative to the remaining techniques. Simultaneously, while the relative performance of Neural Network classifiers improved relative to PLS-DA classifiers, Neural Network classifier performance remained poor compared sPLS-DA, SVM, and Random Forest classifiers. Over the real datasets, a trend of better performance of SVM and Random Forest classifier performance was observed.

## 1. Introduction

As the reactants, intermediates, and products of metabolic reactions, in vivo metabolite concentrations are reflective of stable hereditary factors such as DNA sequence and epigenetic modifications as well as transient stimuli that elicit metabolic responses over varying time domains. Many diseases—including ubiquitous human diseases such as diabetes [1], coronary artery disease [2], heart failure [3], and cancer [4]—are either caused by or result in metabolic dysregulation. Consequently, metabolite concentrations quantified from human samples report both constitutive diseases processes such as atherosclerosis [5] and acute disease events such as myocardial infarction [6] and cerebral infarction [7]. While metabolic phenotyping is well suited to inform clinical phenotype prediction given the interplay between metabolism and human disease, the success of this approach depends on the discriminative power of the statistical classification techniques employed. Consequently, we sought to conduct a thorough and rigorous analysis of classifier techniques for use in metabolomics, with special attention paid to high dimensional data as a common feature of untargeted analyses. In evaluating multiple statistical classification techniques, the maximization of different objective functions will invariably lead to different results. An objective function of maximizing biological knowledge extraction may lead to the choice of simple, human interpretable classifiers. In contrast, an objective function of error minimization may lead to the selection of “black box” classification techniques such as classifier ensembles for which meaningful biological inference cannot be made. In conducting our evaluations, we have defined minimizing classification error and cross-entropy loss objective functions, predicated on the assumption that for metabolite concentrations to inform diagnostic and prognostic predictions in medicine, accuracy is to be valued above model interpretability. In selecting classification techniques to evaluate, we have sought to include classifiers with widespread utilization in metabolomics (e.g. PLS-DA), ensemble methods (e.g. Random Forests), methods that allow non-linear discrimination functions and are robust given non-normal data (e.g. Support Vector Machines and Neural Networks), and methods with embedded feature selection (e.g. Sparse PLS-DA). This work addresses a concern raised by others [8,9] that PLS-DA predominates classification in metabolomics without regard to potential drawbacks or misuses. In order to evaluate classifier performance, we utilized simulation studies designed to emulate an analysis workflow post analytical detection and quantification of metabolite abundances—that is we assume method-specific data processing such as peak detection, signal deconvolution, and chromatographic alignment have already been conducted. While we refer to simulated abundances as metabolites for simplicity our evaluations would generalize to datasets with ion features that have not been grouped or annotated as compounds. In addition to simulation studies, we evaluated classifier performance across two clinical datasets in which a principle aim was using metabolomics to facilitate a diagnostic determination. In the first clinical dataset, DeFilippis et al. [6] sought to use metabolomics to distinguish between subtypes of acute myocardial infarction (MI) and to characterize the disease state changes underlying the transition from stable coronary artery disease (sCAD) to acute MI. In the second dataset, Fahrmann, et al. [10] evaluated biomarkers for the detection of adenocarcinoma lung cancer.

We provide next a brief high-level introduction to the techniques evaluated for phenotype discrimination and an overview of the simulation studies that were designed to be biologically plausible and mimic a metabolomics analysis workflow post metabolite quantification. Partial Least Squares-Discriminant Analysis (PLS-DA) is a ubiquitous classification technique that has been widely utilized in metabolomics studies [9]. The objective of Partial Least Squares (PLS) is to find latent components that maximize the sample covariance between sample phenotype and observed abundance data after applying linear transformations to both [11]. An advantage of PLS approaches is that the latent components are iteratively determined to maximize the remaining phenotype covariance which facilitates straightforward dimension reduction (by considering a parsimonious set of the components that capture sufficient phenotypic variance) and can mitigate estimability issues arising from the presence of more metabolites then samples (*p* > *n*) and from multicollinearity. To generalize PLS regression to classification, a matrix of binary phenotype indicators can be used as dependent variables and a discriminant analysis such as Fisher’s discriminant analysis or nearest centroids can be conducted (hence PLS-DA). Given that metabolomics studies typically have *p* ≫ *n*, that is far more metabolites quantified than samples, variable (metabolite) selection is often advisable. This artifact is especially pronounced when considering data with ion features. Sparse PLS-DA can be conceptualized as a modification of PLS-DA that embeds feature (metabolite) selection through regularization. Sparsity is enforced by penalizing the norm of the weights that define the linear transformations that relate the observed abundance data and the latent components [12,13]. Dependent on the penalization parameter, some of the individual metabolite weights may shrink to zero— effectively removing that metabolite from the model.

While PLS methods aptly handle the multicollinearity present in metabolomics data due to abundance correlations within metabolic pathways, the latent components are linear combinations of the metabolites and assume metabolite abundances are approximately normally distributed. The need for non-linear function approximation is warranted given consideration to the non-linearity of enzyme kinetics (see for example [14]). Support vector machines (SVMs) are binary classifiers that seek to find linear hyperplanes that maximize the separation between classes [15]. SVMs can approximate non-linear decision boundaries between classes by employing a linear or non-linear mapping of the metabolite data to a higher dimensional space in which a separable or nearly-separable linear hyperplane between classes can be found. Since SVMs are binary classifiers, to employ SVMs for multi-phenotype discrimination multiple classifiers need to be constructed and aggregated. In our analyses we employ a “one-against-one” approach [16]. The strength of SVM classifications in non-linear discrimination—of great benefit in metabolomics— stems from the ability of SVMs to approximate arbitrary continuous functions [17] (known as universal approximation). This desirable property has also been shown for Neural Networks (see for example [18] for a proof of universal function approximation for multilayer feedforward networks). Neural Networks are so named as early work in this field [19] focused on developing mathematical models that mimic cognition—specifically recognition via the activation of neurons and propagation of signals. A general feedforward network consists of three types of node layers: an input layer for inputting metabolite abundances, hidden layer(s) conceptualized as neurons that aggregate and process signals, and an output layer that is used for prediction (such as predicting phenotype).

The final classification technique considered in this analysis is Random Forests. A Random Forest is an ensemble of classification or regression trees that employs bootstrap aggregation (“bagging”) and random subspace constraints to minimize the variance of model sampling [20,21]. Bagging is conducted in this context by constructing a collection of individual trees using repeated sampling with replacement from the original data and aggregating the trees into an ensemble for making predictions. Bagging is a form of model averaging that has been shown to increase accuracy in proportion to the degree to which the underlying model is sensitive to perturbations of training data [22]. Random Subspace constraints stipulate that during the iterative process of tree construction, only a random subset of metabolites will be considered in defining branch splits [23]. Enforcing a Random Subspace constraint improves the performance of the bagging strategy by reducing the correlation between the individual trees [21].

We chose to evaluate the performance of the selected classification techniques using simulated datasets as evaluating performance on a single or small collection of real datasets would exhibit high variance. Evaluating performance on a large number of simulated datasets allows for more precise estimates of relative performance and for directly evaluating the effects of increased noise, increased non-linearity, and/or departures from approximate normality. A significant hurdle in simulating metabolomics data is that such data is marked by a significant degree of pairwise and higher order partial correlations [24,25]. Metabolites in the same reaction or linked reactions function as substrates, intermediates, and products thus generating complex correlation structures. As a result, simulating metabolomics data necessitates generating multivariate distributions of correlated metabolites. Further, as the enzymes that catalyze biochemical reactions are often subject to regulatory processes such as feedback inhibition [14], complex partial correlation structures must also be simulated. Acknowledging this, we simulated metabolite data in blocks representing biological processes. To ensure diversity in the simulation studies, random correlation matrices were generated for each simulation. Generating random correlation matrices requires specialized methods to ensure that the resulting matrices are positive definite. For this we employed the method developed by Lewandowski, et al. [26] which generates partial correlations using a graph (network) structure known as a C-vine. Further details are contained in the Methods section.

## 2. Results

### 2.1. Simulated metabolomics data

For both the baseline and non-linear scenarios, 1,000 simulation studies were conducted. The C- vine procedure (Figure 1) for generating random covariance matrices facilitated generating clusters of simulated metabolites to mimic discrete biological processes. An important aspect of this is the generation of partial correlations as regulatory mechanisms such as feedback inhibition may generate such relationships.

**Figure 1.**
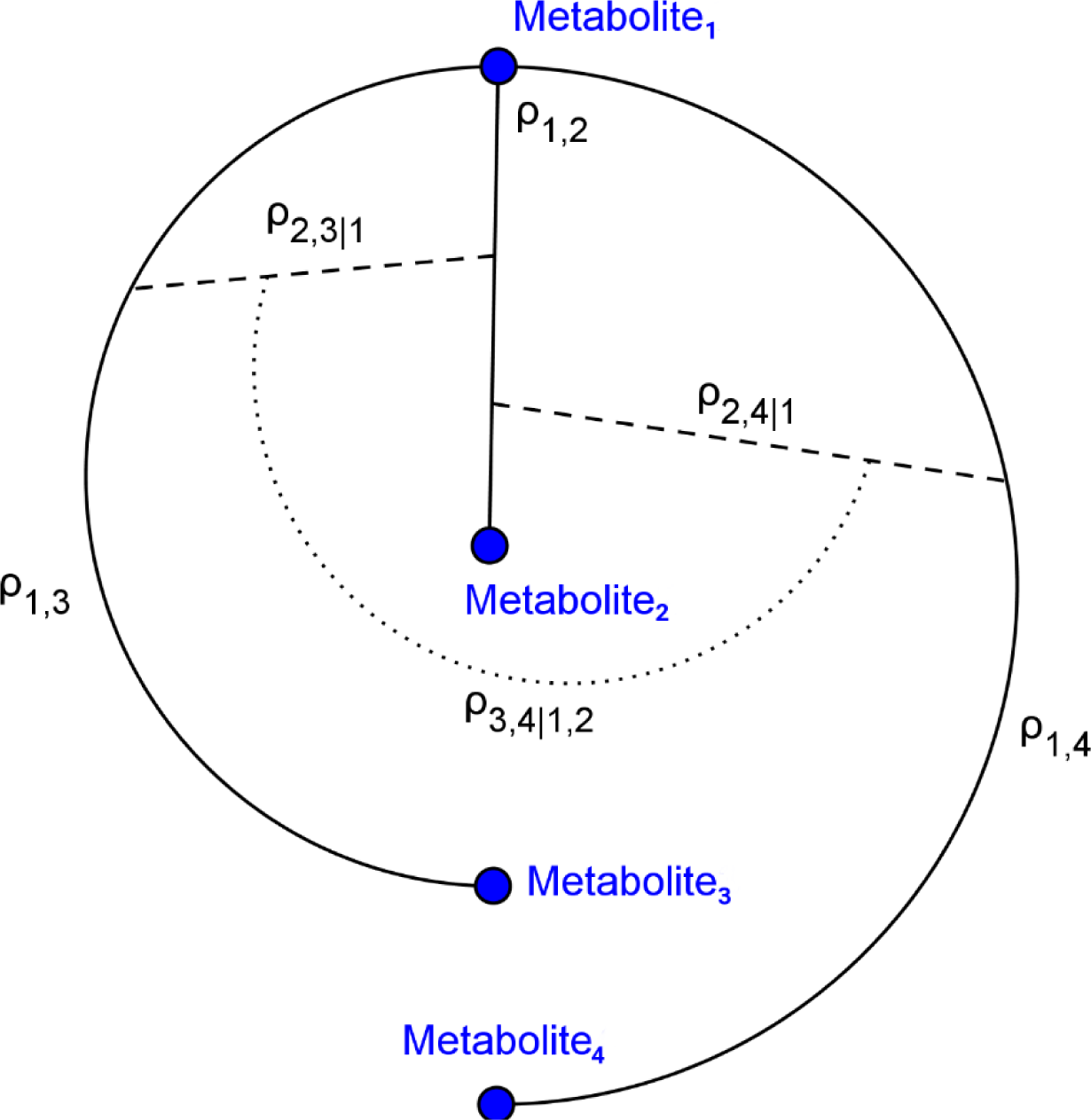
C-vine graph illustrating partial correlation structure. C-vines were utilized to generate biologically plausible metabolomics data. *p_i,j_* represents the correlation between metabolites *i* and *j*. *p_i,j|k_* represents the partial correlation between metabolites *i* and *j* after conditioning on *p_k,i_* and *p_k,j_*.

Figure 2 depicts the simulated metabolite abundance data from a randomly selected baseline scenario study both prior-to and post-significance filtering. Blocks of correlated metabolites are visible (column clusters) as expected given the data generation procedure. In the baseline scenarios classifiers were evaluated on both the prior-to and post-significance filtered datasets such as those shown in Figure 2; in the non-linear scenarios, further transformation was conducted. Figure 3 illustrates how abundance data was generated to follow a variety of random non-normal distributions for the non-linear scenarios. Throughout the non- linear scenario data generation process, blocks of metabolites were simulated to follow a multivariate Gaussian distribution and a random non-linear transformation was applied over each block.

**Figure 2.**
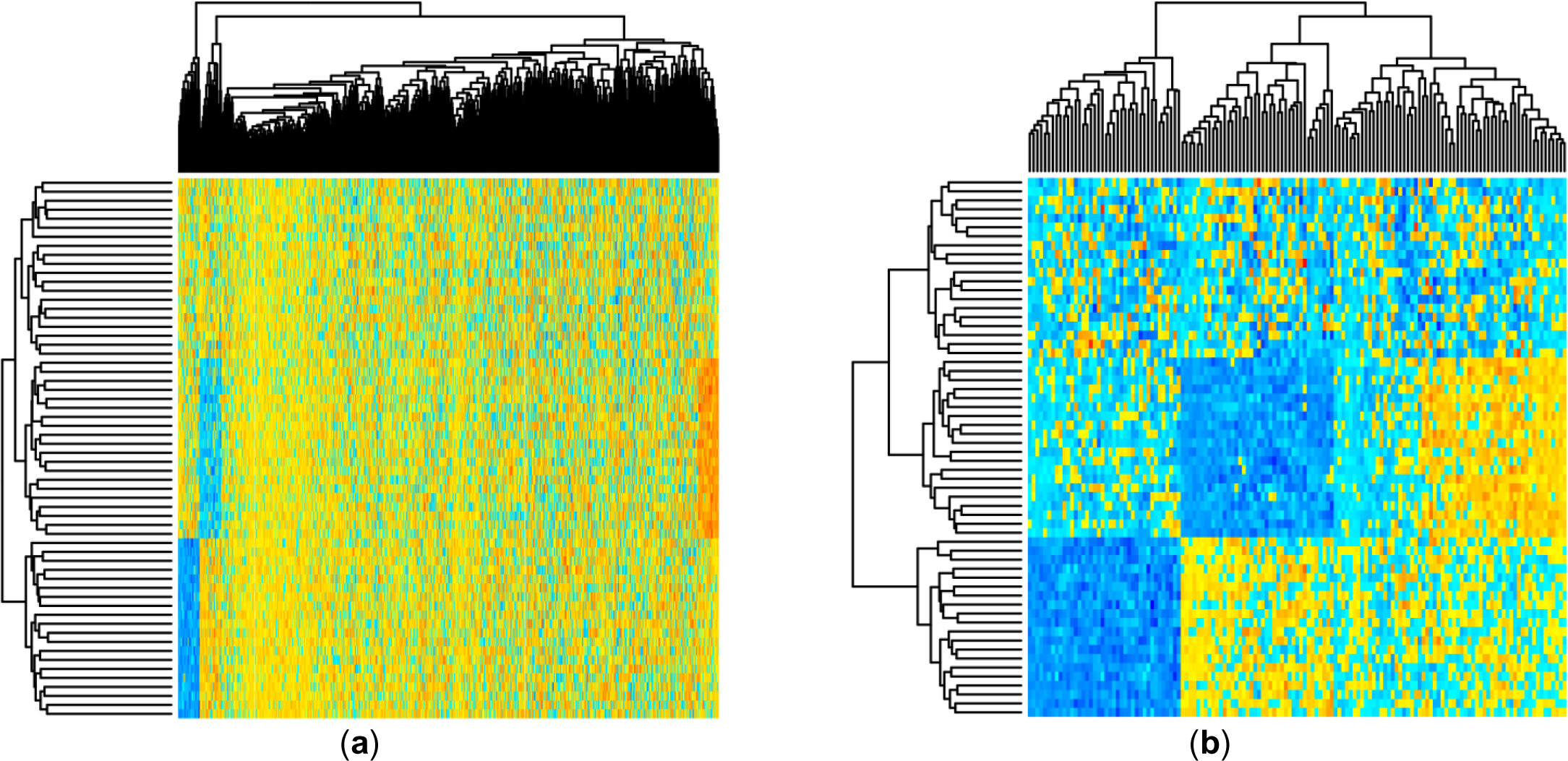
Heatmap showing simulated metabolite abundance data from a randomly selected baseline scenario before and after significance filtering. (**a**) Prior to significance filtering: Distinct clusters of metabolites can be discriminated as expected given the block-wise generation of correlated metabolites. (**b**) Post-significance filtering.

**Figure 3.**
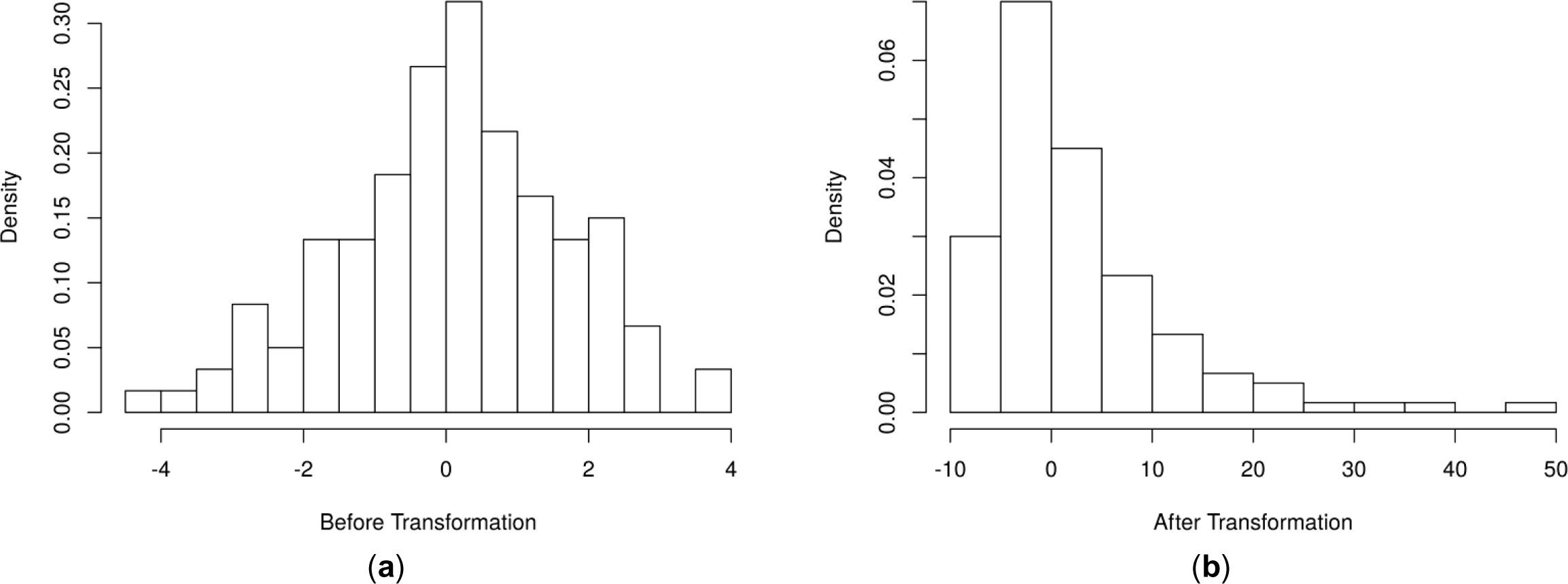
Histogram of an example simulated metabolite abundance distributions for each scenario. (**a**) Baseline scenarios: metabolite abundances were simulated from multivariate normal distributions representing discrete biological processes (one metabolite shown). (**b**) Non-linear scenarios: metabolite abundances were first simulated as in (a) and then random non-linear transformations were applied block-wise.

### 2.2. Evaluation of classifier performance in simulation studies

#### 2.2.1. Aggregate performance

The misclassification rate for each technique over the simulation studies are summarized in Table 1. Over the baseline scenarios and prior-to significance filtering, sPLS-DA exhibited a lower misclassification rate than the remaining techniques (Median ± IQR: 3.3% ± 21.7%). The performance of PLS-DA and Random Forests was similar with respect to median misclassification rate. Neural Networks had a higher median misclassification rate than the other techniques, while SVMs demonstrated the greatest spread (by SD or IQR). The application of significance filtering improved the mean and median misclassification rate for each technique. In the non-linear scenarios (prior-to and post-significance filtering), the performance of PLS-DA deteriorated significantly more than the other techniques. Post- significance filtering, sPLS-DA and SVM classifiers exhibited similar median misclassification rates that were lower (better) than Random Forests and Neural Networks (in that order).

**Table 1.** Misclassification rate (%) observed by technique and by significance filtering status (pre- vs. post) throughout the 1,000 baseline simulation studies and 1,000 non-linear simulation studies. The lowest median misclassification rate observed over each scenario type is shown in bold face.

| Method | Baseline Pre- |  | Baseline Post- |  | Non-linear Pre- |  | Non-linear Post- |  |
| --- | --- | --- | --- | --- | --- | --- | --- | --- |
|  | Mean±SD | Median±IQR | Mean±SD | Median±IQR | Mean±SD | Median±IQR | Mean±SD | Median±IQR |
| PLS-DA | 15.1±15.5 | 10.0±26.7 | 12.9±15.4 | 5.0±25.0 | 35.1±19.2 | 35.0±28.3 | 21.8±17.9 | 21.7±28.3 |
| sPLS-DA | 11.3±14.5 | **3.3±21.7** | 10.5±14.2 | 1.7±20.0 | 12.2±15.4 | **3.3±25.0** | 11.9±15.5 | 3.3±25.0 |
| SVM | 22.0±23.5 | 10.0±40.0 | 10.0±14.1 | **1.7±18.3** | 24.2±25.3 | 13.3±41.7 | 11.8±15.2 | **3.3±23.3** |
| NNet | 21.9±15.9 | 20.0±25.0 | 15.6±13.8 | 10.0±20.4 | 28.6±17.6 | 28.3±26.7 | 16.5±15.7 | 11.7±25.0 |
| RF | 14.9±14.7 | 10.0±25.0 | 12.9±14.1 | 8.3±23.3 | 14.5±15.0 | 8.3±25.0 | 13.4±15.1 | 6.7±23.3 |

Cross-entropy loss over the simulation studies is summarize in Table 2. Over the baseline scenarios prior-to significance filtering, SVM and Random Forest classifiers exhibited similar performance (Median ± IQR: 0.56 ± 0.50 and 0.70 ± 0.64, respectively); PLS-DA, sPLS-DA, and Neural Networks were similar and higher than SVM/RF classifiers. Post-significance filtering in the baseline scenarios, SVM classifiers exhibited the lowest cross-entropy loss, followed by Random Forests. As before, PLS-DA, sPLS-DA, and Neural Networks showed similar performance. In the non-linear scenarios, the qualitative pattern of results was similar to the baseline scenarios (Table 2).

**Table 2.** Cross-entropy loss observed by technique and by significance filtering status (pre- vs. post) throughout the 1,000 baseline simulation studies and 1,000 non-linear simulation studies. The lowest median cross-entropy loss observed over each scenario type is shown in bold face.

| Method | Baseline Pre- |  | Baseline Post- |  | Non-linear Pre- |  | Non-linear Post- |  |
| --- | --- | --- | --- | --- | --- | --- | --- | --- |
|  | Mean±SD | Median±IQR | Mean±SD | Median±IQR | Mean±SD | Median±IQR | Mean±SD | Median±IQR |
| PLS-DA | 1.19±0.15 | 1.20±0.22 | 1.08±0.17 | 1.06±0.27 | 1.36±0.16 | 1.36±0.24 | 1.17±0.19 | 1.18±0.26 |
| sPLS-DA | 1.11±0.14 | 1.08±0.19 | 1.05±0.16 | 1.02±0.25 | 1.11±0.15 | 1.08±0.20 | 1.05±0.17 | 1.01±0.28 |
| SVM | 0.63±0.45 | 0.70±0.64 | 0.43±0.37 | **0.23±0.49** | 0.69±0.47 | 0.71±0.64 | 0.48±0.39 | **0.30±0.55** |
| NNet | 1.12±0.20 | 1.10±0.33 | 1.02±0.18 | 0.96±0.29 | 1.20±0.23 | 1.19±0.34 | 1.03±0.21 | 0.97±0.34 |
| RF | 0.58±0.35 | **0.56±0.50** | 0.55±0.33 | 0.51±0.50 | 0.57±0.36 | **0.52±0.52** | 0.54±0.35 | 0.48±0.50 |

#### 2.2.2. Performance within simulation studies

Pairwise comparisons of misclassification rate within the simulation studies are shown in Figure 6. For example, PLS-DA had a lower misclassification rate than sPLS-DA in 11.00% of the 1,000 baseline scenario studies prior-to significance filtering (Figure 5(a), row 1, column 2) and in 15.21% of the studies post-significance filtering (Figure 5(b) row 1, column 2). PLS-DA exhibited greater misclassification rate relative to the other techniques in the majority of studies with the exception of Neural Networks given baseline scenarios. In the majority of non-linear scenario studies, Neural Networks exhibited lower misclassification rates than PLS-DA. Overall, sPLS-DA had a lower misclassification rate than each other technique more often than not with the exception of SVM classifiers in the non-linear scenarios prior-to significance filtering.

**Figure 4.**
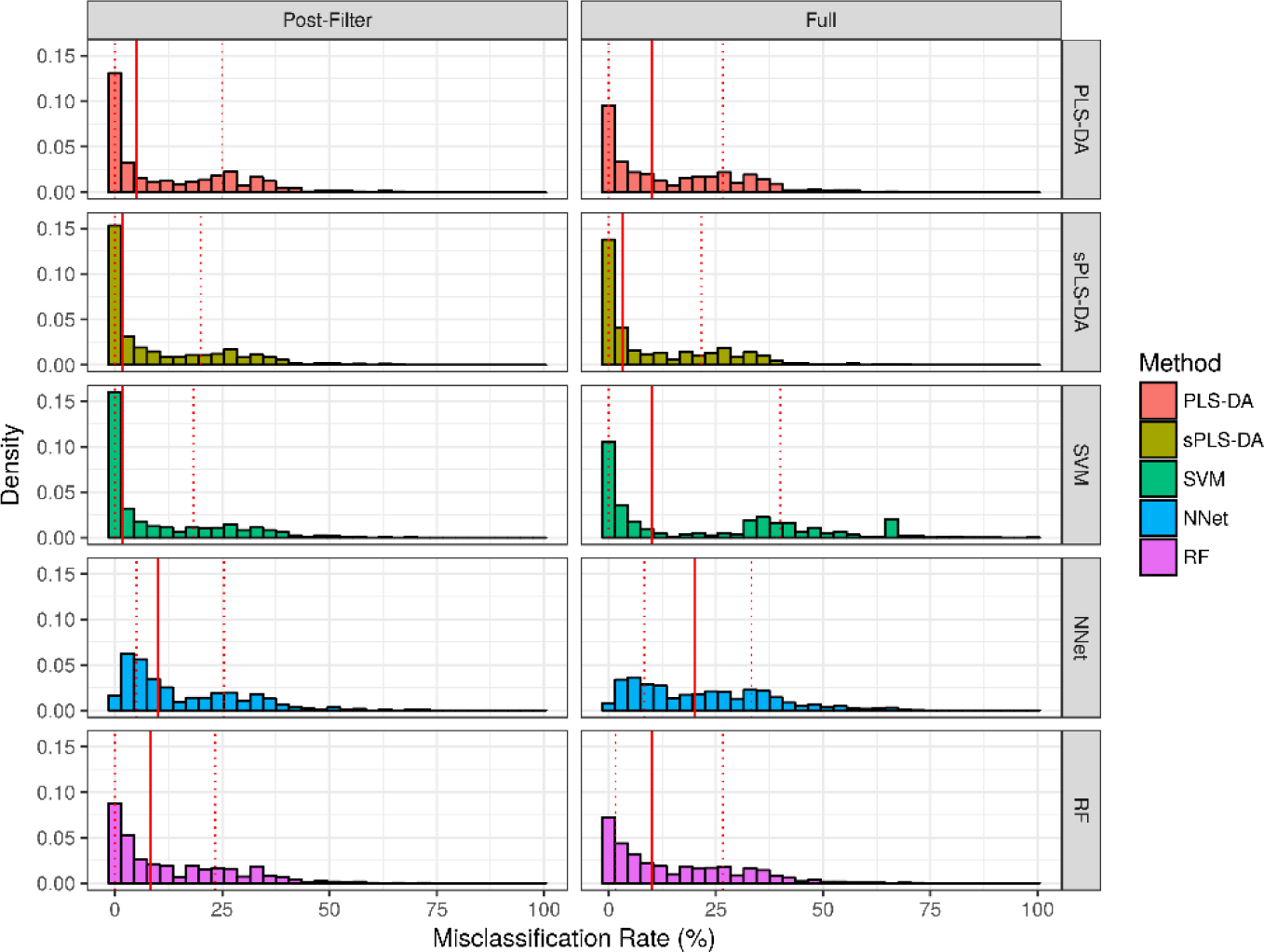
Misclassification rate observed in baseline scenario simulation studies. Solid red line represents the median of each distribution, while dashed red lines represent the 1^st^ and 3^rd^ quantiles (25^th^ and 75^th^ percentile).

**Figure 5.**
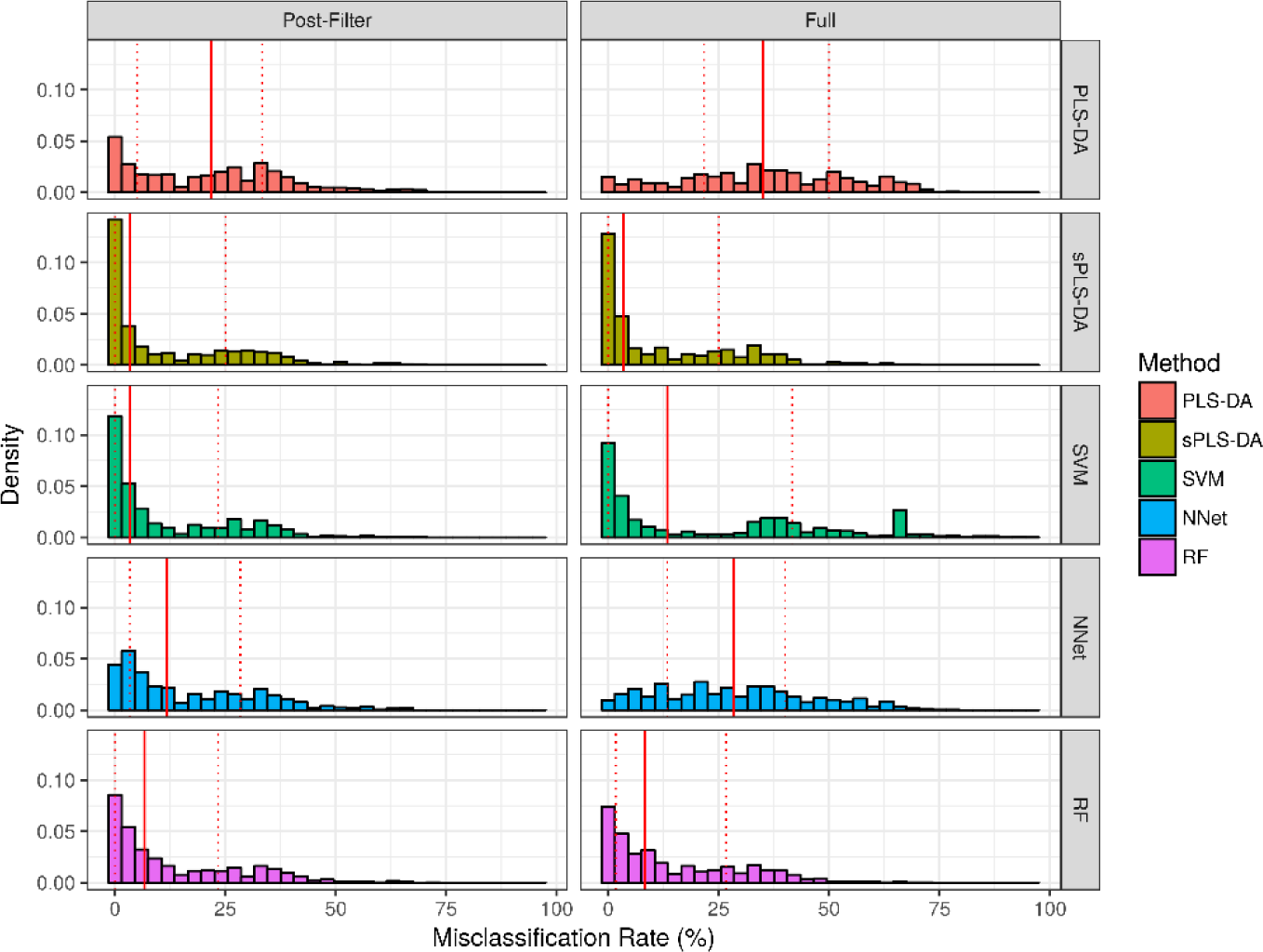
Misclassification rate observed in the non-linear scenario simulation studies. Solid red line represents the median of each distribution, while dashed red lines represent the 1st and 3rd quantiles (25th and 75th percentile).

**Figure 6.**
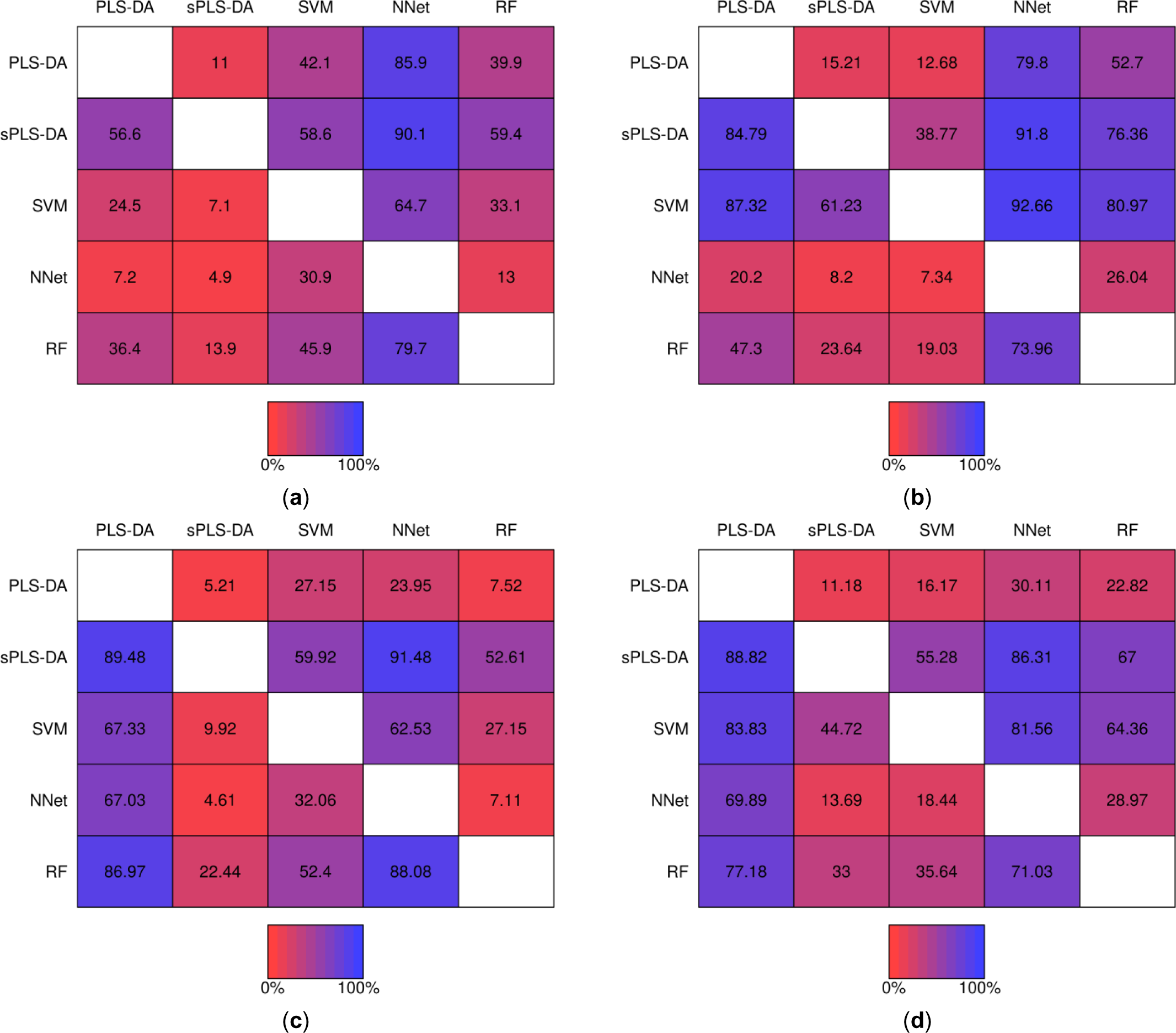
Matrices showing the proportion of the time a fixed technique performed better than another fixed technique during the same simulation study. Proportions were computed from the 1,000 baseline simulation studies prior-to significance filtering (**a**) and post-significance filtering (**b**); and from the 1,000 non-linear simulation studies prior-to significance filtering (**c**) and post-significance filtering (**d**).

### 2.3. Performance over real datasets

Performance over the real datasets is shown in Table 3. Over the Adenocarcinoma study data, Neural Networks observed the lowest misclassification rate (14.3%) on the test dataset prior to significance filtering with SVM classifiers demonstrating lower misclassification (10.7%) over the test data post-significance filtering. With respect to cross-entropy loss the same was observed, while Random Forests also demonstrated equally low cross-entropy loss as Neural Networks prior to filtering. Over the acute MI study data, Random Forest classifiers had the lowest misclassification rate estimated by cross-validation prior-to and post- significance filtering (28.9% and 10.5%). With respect to cross-entropy loss, Random Forest classifiers demonstrated lowest cross-validation estimated loss, while SVM classifiers demonstrated the lowest loss following significance filtering.

**Table 3.** Misclassification rate and cross-entropy loss observed over real datasets.

| Dataset | Technique | Mis. (%) |  | Cross-Entropy Loss |  |
| --- | --- | --- | --- | --- | --- |
|  |  | Pre- | Post- | Pre- | Post- |
| Adenocarcinoma | PLS-DA | 17.9 | 17.9 | 0.78 | 0.75 |
|  | sPLS-DA | 32.1 | 14.3 | 0.83 | 0.72 |
|  | RF | 17.9 | 14.3 | **0.68** | 0.57 |
|  | SVM | 21.4 | **10.7** | 0.78 | **0.53** |
|  | NNet | **14.3** | 21.4 | **0.68** | 0.77 |
| Acute MI | PLS-DA | 47.4 | 50.0 | 1.52 | 1.48 |
|  | sPLS-DA | 34.2 | 18.4 | 1.42 | 1.16 |
|  | RF | **28.9** | **10.5** | **1.10** | 0.71 |
|  | SVM | 50.0 | 15.8 | 1.55 | **0.66** |
|  | NNet | 47.4 | 31.6 | 1.47 | 1.24 |

## 3. Discussion

In this report, we have detailed a rigorous and comprehensive evaluation of selected statistical classification techniques for discrimination of phenotype given metabolomic data. While at least one ad-hoc analysis of classifier performance has been conducted on a specific dataset [8], an analysis utilizing realistic and diverse simulation studies to generate consistent estimates of performance had not been conducted previously. By varying many parameters of the simulation studies including the number of metabolite clusters that differ between phenotypes, the effect size of differences, and the degree of departure from approximate normality we have ensured that estimates of classifier performance are sufficiently general.

A few key conclusions are supported by the analysis of misclassification rate. First, the performance of PLS-DA and Neural Network classifiers was generally worse than other classification techniques. The performance deficit of PLS-DA compared to other techniques was especially pronounced with the introduction of non-normal error distributions for metabolite clusters. Interestingly, within the same simulation studies, Sparse PLS-DA classifiers exhibited a lower misclassification rate than each other technique with one scenario exception (SVMs performed better post-significance filtering given non-linear data). This suggests that regularization of PLS-DA improves performance in addition to encouraging a more sparse classifier. The performance deficit of Neural Network classifiers may be attributed to model complexity. For example, the number of parameters (weights) estimated for an Neural Network classifier with one hidden layer of 10 nodes given three phenotypes and 1,000 metabolite features is 10,058 (with bias nodes). Contrastingly, a PLS-DA model with two latent components would have 2,000 loading weights in addition to regression coefficients.

With respect to cross-entropy loss, Random Forests exhibited the lowest median loss prior-to significance filtering while SVM classifiers exhibited the lowest median loss post-significance filtering. It is interesting to note that while sPLS-DA classifiers performed generally well with respect to misclassification rate in comparison to the remaining techniques, this was not observed using cross-entropy loss as an objective function for measuring performance. This shows that choice of loss function has an impact on the measurement of relative classifier performance. While an extensive discussion of the relative merits of different loss functions are beyond the scope of the current work, we do postulate that there may be situations where the cross-entropy loss is more appropriate than the 0-1 loss that yields the misclassification rate as the empirical risk.

A limitation of this work is that while we have sought to minimize the effect of algorithm parameters on observed misclassification rate and cross-entropy loss by conducting extensive parameter tuning via cross-validation, the entire parameter space was not evaluated for multiple techniques. For example, while a thorough grid search (with smoothing) was conducted to select the Gaussian kernel bandwidth parameter for the SVM classifiers, the space of kernels not evaluated remains infinite. Additionally, in the current study we have defined measures of prediction error as the objective criteria for measuring classifier performance. However, other criteria such as model interpretability may be important for practitioners. This is especially the case when classifier techniques are used for hypothesis testing or for biological inference.

## 4. Materials and Methods

### 4.1. Simulated metabolomics data

Evaluation of classifier techniques for metabolomics-based phenotype discrimination requires simulation studies that realistically mimic data captured using analytical methods such as nuclear magnetic resonance or chromatography-coupled mass spectrometry from biological samples (e.g. cell or biofluid extract). While the distribution of metabolite abundances may have platform and/or sample medium specific artifacts, we posit that four features are common to untargeted metabolomics studies: (1) significant correlations and higher-order partial correlations between metabolites within biological processes, (2) a small proportion of differentially abundant metabolites localized specific biological processes, (3) a large number of quantified metabolites relative to sample size-most demonstrating variance orthogonal to phenotype, and (4) non-Gaussian error distributions and non-linear relationships between metabolite abundances and phenotype attributes.

Metabolites within biochemical processes are related by substrate, intermediate, and product relations thus generating complex correlation structures. Consequently, we generated simulated abundance data to follow multivariate distributions with covariance structures that allow for mimicking biological processes. Further, as the enzymes that catalyze biochemical reactions are often subject to regulatory processes such as feedback inhibition [14], we simulated complex partial correlation structures. We represent metabolite abundance data as a matrix ***X*** of dimension ***n** × **p*** given ***n*** samples and ***p*** metabolites, sample phenotype labels as a vector ***y*** or as a matrix of binary indicators ***Y***. For each simulation study dataset we generated 40 multivariate blocks of 25 metabolites. Each block, ***X**_k_* was generated such that ***X**_k_* followed a multivariate Gaussian distribution, that is: ***X**_k_ ~ N* (***m**_k_*, ***Ʃ**_k_*. The covariance matrices ***Ʃ**_k_* were each randomly generated using C-vines for simulating partial correlations between metabolites [26]. The algorithm for generating correlation matrices utilizing C-vines is presented below and was developed by Lewandowski, Kurowicka and Joe [26]:

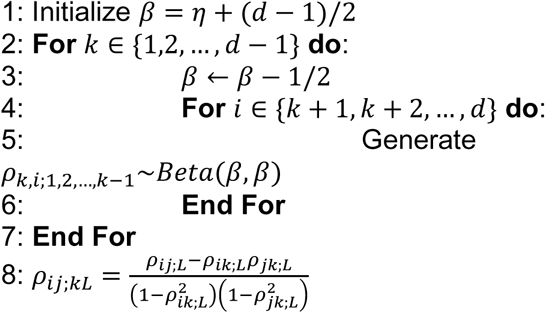

Three phenotypes were simulated by supplying different means for a small proportion of simulated metabolite blocks. A reference phenotype had ***m**_k_* = 0 for all *k*. The number of perturbed blocks in the comparator phenotypes was generated to follow a discrete uniform distribution, *Unif*(1,5). The perturbed block means were generated using a hierarchical model with *m_k_ ~ N* (***θ**_k_*,1) and *θ_k_ ~ Exp* (1/2). A simulated Bernoulli process with *p* = 1/2 was employed to modulate the sign of *θ_k_*. The “non-linear” scenario data was generated as above with the added data generation step of applying a non-linear transformation to the empirical cumulative distribution function of the multivariate Gaussian blocks to generate randomly-parameterized General Gaussian Distributions (GGD). The probability distribution function of a GGD with location parameter zero is defined as [27]:

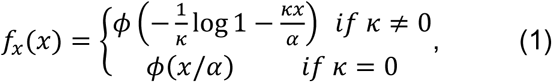

where *ϕ* is the standard Gaussian probability distribution function.

To evaluate the hypothesis that significance filtering prior to classifier construction would have an impact on the relative performances of the techniques evaluated, within each simulation study we evaluated performance prior-to and post- significance filtering. Significance filtering was conducted by filtering on metabolites with significant pairwise *t*-tests between groups at a significance level of *α* = 0.025.

### 4.2. Classification techniques

#### 4.2.1. Partial Least Squares-Discriminant Analysis (PLS-DA)

Partial least squares (for linear regression: PLS-R) has the following model formulation [28-30]:

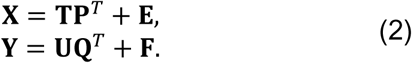

In this formulation, **E** and **F** represent Gaussian noise, **T** and **U** are latent component matrices, and **P** and **Q** are the loading matrices that relate the latent components to the observed metabolite abundances **X** and the observed response variables **Y**. PLS algorithms seek to find weight vectors **w** and **c** such that [Cov(**t**, **u**)]^2^ = [Cov(**Xw**, **Yc**)]^2^ is maximized [31]. Specifically, the Nonlinear Iterative Partial Least Squares Algorithm (NIPALS) may be used to find **w** and **c**. The algorithm pseudocode is presented as in Rosipal [32] below. Until convergence, repeat:

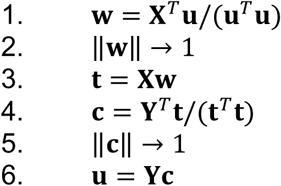

Finally, **p** and **q** can be found by ordinary least squares (OLS) regression and **X** and **Y** are deflated. In our simulation studies and applications, we consider the first three score vectors, 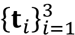. To utilize PLS for classification **Y** is defined as a binary indicator matrix of sample phenotypes and a discriminant analysis is conducted following regression.

#### 4.2.2. Sparse Partial Least Squares-Discriminant Analysis (sPLS-DA)

Lê Cao, *et al.* [33] proposed a *l*_1_ regularized version of PLS-DA to encourage sparsity in PLS modeling. In this section we modify the description found in Lê Cao, *et al.* [34] to maintain consistency. The objective of sPLS remains to find **w** and **c** such that [Cov(**Xw**,**Yc**)]^2^ is maximized, but now subject to penalization of the norm of **w** To proceed we introduce a result from Höskuldsson [31], that **w** and **c** are the vectors that satisfy:

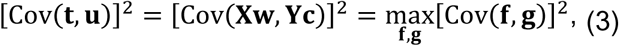

given the singular value decomposition: 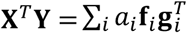 Consequently, the regularized optimization problem can be restated as:

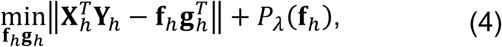

where *h* = 1,2, … , *H* is the number of deflations. The penalization parameter λ was selected in each simulation study or real data analysis utilizing a grid search strategy (see Section 4.3).

#### 4.2.3. Support Vector Machines (SVM)

Support vector machines (SVM) are binary classifiers that seek to find hyperplanes that maximize the separation between classes. These hyperplanes may be linear in the original space of metabolite abundances (of dimension *p*) or in a higher dimensional space (of dimension *p*′) that allow for non-linear boundaries in the original space [21]. A decision hyperplane for binary classification with 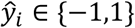 as phenotype indicators has the following form [15]:

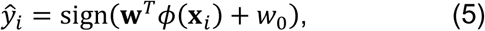

where *ϕ* is an arbitrary real valued function and **w** is a vector of weights. This leads to the following optimization problem for *M* = 1/||**w**|| [15,21]:

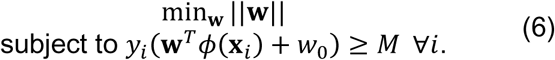

As the optimization problem in (6) seeks to maximize the margin, *M*, that separates the phenotypes, SVM classifiers are often referred to as maximal margin classifiers. In the case that a hyperplane does not separate the observations by phenotype, then slack terms, *ξ_i_*, are added allowing for a “soft” margin and yielding the optimization problem:

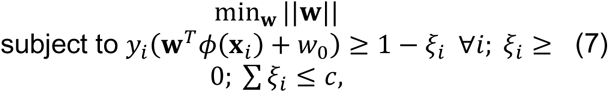

where *c* is a constant. The optimization problem is then solved by quadratic programming utilizing Lagrange multipliers. Conveniently, this optimization does not require explicit computation of the original data given new basis functions, that is *ϕ*(***x**_i_*), but rather the inner products 〈*ϕ*(***x**_i_*), *ϕ*(***x**_i_*′)〉 [15,21]. Consequently, non-linear transformations are usually defined in terms of the kernel function determined by the inner product, *K*(*x*, *x*′) = 〈*ϕ*(***x**_i_*), *ϕ*(***x**_i_*′)〉. In each simulation study or real data analysis, Radial (Gaussian) kernels: *K*(*x*, *x*′) = exp(−-γ||*x* - *x*′ ||^2^), were utilized with γ selected using a grid search strategy (see Section 4.3).

#### 4.2.4. Neural Networks (NNet)

In this analysis, we evaluated feed-forward neural networks for classification. A feed forward network is a class of directed acyclic graphs loosely inspired by models of cognition in which metabolite abundances are conceptualized as stimuli and phenotype predictions are conceptualized as perceptions [15,35]. Topologically, a feed-forward network consists of an input layer (allowing for the transfer of metabolite abundances), one or more hidden layers for processing and aggregation of signals from earlier layers, and an output layer with each phenotype represented by a node. Bias nodes may be incorporated to introduce signal independent of topologically antecedent layers. Given this topological representation, the general formula (output for each phenotype) with implicit bias terms is [35]:

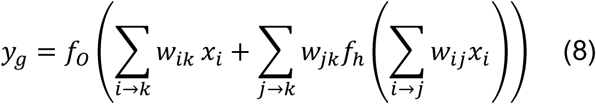

where *f_o_* and *f_h_* are continuous functions applied at output and hidden layer vertices, respectively; *i* → *j* represent directed edges between input layer vertices and hidden layer vertices; *j* → *k* represent directed edges between hidden layer vertices and output layer vertices; and *i* → *k* represent skip-layer transfers from input layer vertices directly to output layer vertices. In our analyses we have utilized “Resilient Backpropagation” (RPROP) for training Neural Network classifiers [36]. In general, backpropagation algorithms iteratively use training observations to compute the output of a network (“forward pass”) followed by computation of the partial derivatives of the error function with respect to network weights (“backward pass”) for updating the weights by gradient descent [35,36]. Resilient backpropagation modifies the weight updating step to adaptively modulate the magnitude of weight updating based on the sign of the partial derivatives [36].

#### 4.2.5 Random Forests

A Random Forest (RF) classifier can be conceptualized as an ensemble of γ classification trees each constructed utilizing a bootstrap sample from the original data. The process of constructing individual classification trees proceeds by recursive binary splits (splitting a parent node into two daughter nodes) selected from a restricted subset of random variables (metabolites) and cut-points [20,21]. Specifically, at each iteration, a set of candidate regions:

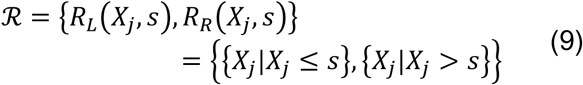

is generated following the selection of a set of random variables (metabolites) sampled with replacement from the bootstrapped data. After generating the regions, the empirical phenotype distribution is computed over each region *R*, that is:

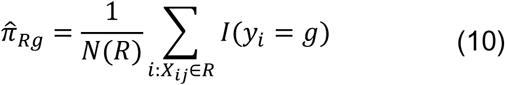

for each phenotype *g* For each region a phenotype is then ascribed:

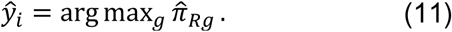

*X_j_* and *s* are then chosen to minimze a measure of node impurity, in our case the misclassification error: 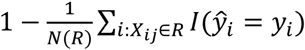 Once *X_j_* and *s* have been selected the current parent node is split into the daughter nodes satisfying {*x_i_*: *x_ij_* ≤ *s*} or {*x_i_*: *x_ij_* > *s*}. When generating an ensemble of individual classification trees, the correlation between individual trees estimated from the bootstrapped samples is reduced by enforcing a random subspaces constraint [23], considering at each binary split only a randomly drawn subset of variables (metabolites). Once an ensemble of trees has been aggregated as a Random Forest, predicted phenotype probabilities can be determined by aggregating individual tree predictions.

### 4.3. Parameter selection

Each of the classification techniques evaluated in the present study represent families of classifiers whose members are uniquely determined by algorithm parameters. Consequently, we sought to minimize the probability that an observed relative difference in classifier performance was due to sub- optimal parameter selection for one or more techniques. During the course of each simulation study (prior-to and post-significance filtering), parameter selection was conducted by minimizing expected cross-entropy loss estimated by cross-validation and smoothed over a parameter grid using kernel smoothing. For reproducibility, the relevant fixed and cross-validation selected algorithm parameters used in defining the classifiers are shown in Table 4.

**Table 4.**
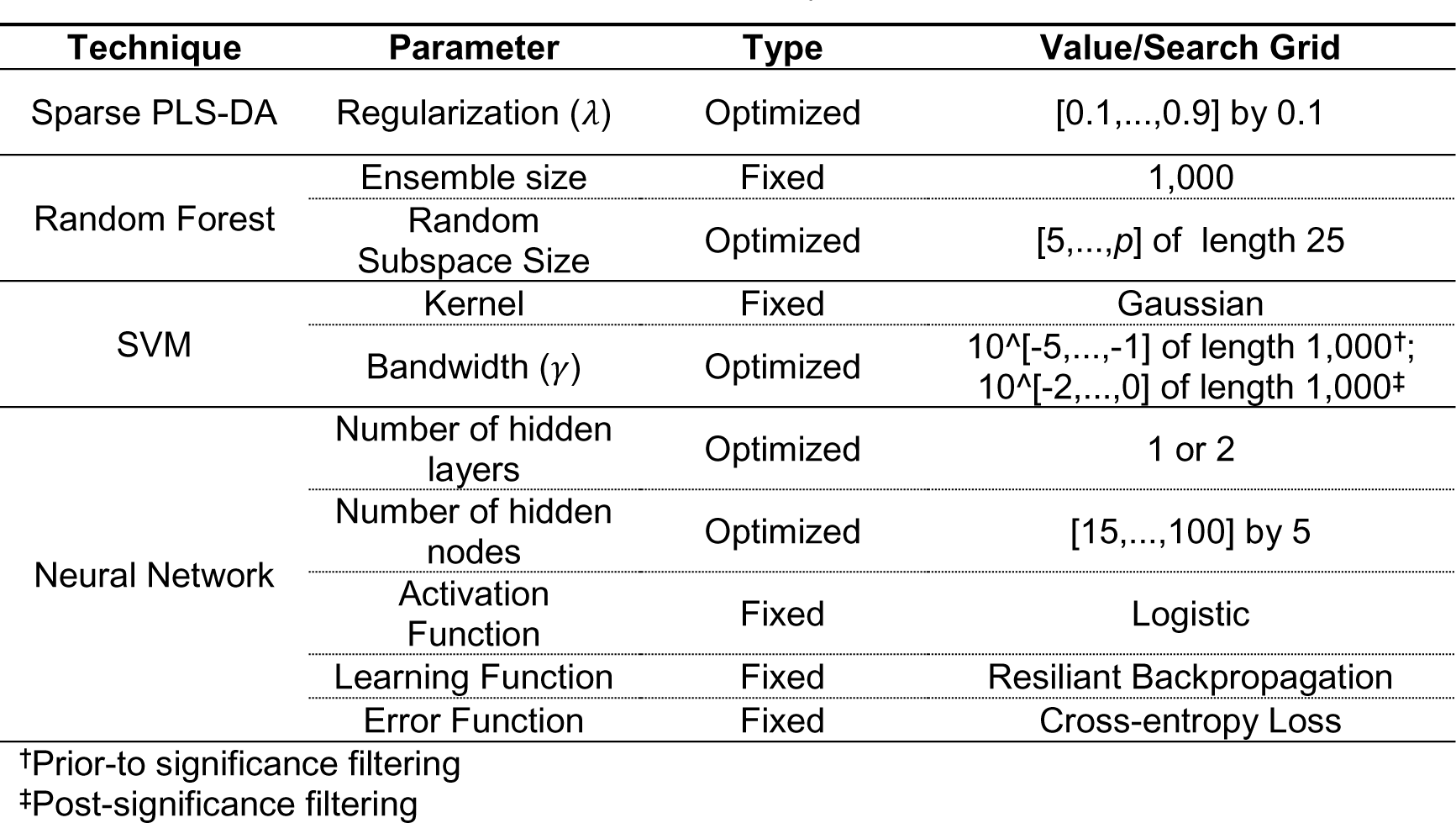
Simulation study parameters

### 4.4. Evaluation of classifier performance

Classifier performance was evaluated by computation of the empirical risk (error) associated with two different loss functions [37]. Defining a phenotype prediction 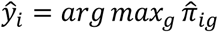 from a classifier, a 0-1 loss function is: 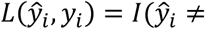 *y_i_*), with associated empirical risk (the misclassification rate): 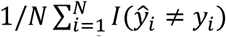. Cross-entropy loss is defined as: 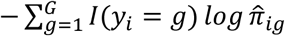 with empirical error: 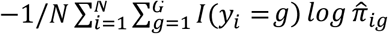 While the misclassification rate measures the frequency of a classifier incorrectly classifying observations, the empirical cross-entropy error measures the average amount of extra information required to represent the true phenotypes with the predicted phenotypes. Consequently, the empirical cross-entropy error provides a measure of how well the predicted phenotypes the true phenotypes. The distinction between these loss functions can be observed with the following case. Given a binary classification task, a misclassified observation with a predicted phenotype probability of 49% incurs less cross-entropy loss than a predicted phenotype probability of 0.1%. Given a 0-1 loss function the computed loss would be the same for a misclassified observation with a predicted phenotype probability of 49% as a predicted phenotype probability of 0.1%.

### 4.5. Clinical datasets

In addition to evaluation of classifier performance via simulation studies, classifier performance was evaluated over two clinical datasets. In the first, DeFilippis et al. [6] employed an untargeted approach for determining a plasma signature that differentiates between thrombotic myocardial infarction (MI), non-thrombotic MI, and stable coronary artery disease (CAD). Thrombotic MI is characterized by atherosclerotic plaque rupture/disruption that leads to the formation of a thrombus and the obstruction of a coronary artery [38] while non-thrombotic MI occur secondary to other causes such as blood supply demand mismatch during tachyarrhythmias, coronary artery spasm or low blood oxygen levels. Plasma samples from 23 subjects presenting with acute MI and 15 subjects with stable coronary artery disease undergoing cardiac catheterization were analyzed. Of the 23 acute MI subjects, 11 were adjudicated to be thrombotic MI and 12 were adjudicated to be non-thrombotic MI utilizing a strict criteria. 1,032 metabolites were detected and quantified by gas chromatography mass spectrometry (GC-MS with electron ionization), and ultra performance liquid chromatography mass spectrometry (UPLC-MS with electrospray ionization) in both positive and negative ion modes. Given the limited sample size we employed a cross-validation approach to measuring classifier performance. In the second dataset, Fahrman et al. [10] sought to determine plasma or serum based biomarkers that could be used to detect adenocarcinoma lung cancer with better specificity than existing methods such as low-dose computed tomography. The researchers developed two case-control cohorts for the purpose of discovering and validating biomarkers of adenocarcinoma lung cancer. Untargeted gas chromatography time-of-flight mass spectrometry with electron ionization was used to determine metabolic abundances in both the discovery and validation cohorts. In our analysis of classifier performance, we utilized the plasma sample metabolite abundances from the second cohort and employed a train-test approach. In the second cohort abundances of 413 metabolites were reported.

## Supplementary Materials

The following are available online at www.mdpi.com/link, Figure S1: title, Table S1: title, Video S1: title.

## Acknowledgments

This work was supported in part by a grant from the American Heart Association (11CRP7300003) and the National Institute of General Medical Sciences (GM103492). Dr. Rai was supported by Wendell Cherry Chair in Clinical Trial Research and generous support from the James Graham Brown Cancer Center.

The authors thank Samantha M. Carlisle for her review and insights.

## Author Contributions

A.P.D. and S.N.R. conceived and designed the experiments; P.J.T. performed the experiments and analyzed the data; P.J.T. wrote the paper.

## References

1. Del Prato, S.; Marchetti, P.; Bonadonna, R.C. Phasic insulin release and metabolic regulation in type 2 diabetes. Diabetes 2002, 51 Suppl 1, S109–116.

2. Freeman, M.W. Lipid metabolism and coronary artery disease. 2006, 130–137.

3. Ashrafian, H.; Frenneaux, M.P.; Opie, L.H. Metabolic mechanisms in heart failure. Circulation 2007, 116, 434–448.

4. Cairns, R.A.; Harris, I.S.; Mak, T.W. Regulation of cancer cell metabolism. Nat Rev Cancer 2011, 11, 85–95.

5. Chen, X.; Liu, L.; Palacios, G.; Gao, J.; Zhang, N.; Li, G.; Lu, J.; Song, T.; Zhang, Y.; Lv, H. Plasma metabolomics reveals biomarkers of the atherosclerosis. Journal of Separation Science 2010, 33, 2776–2783.

6. DeFilippis, A.P.; Trainor, P.J.; Hill, B.G.; Amraotkar, A.R.; Rai, S.N.; Hirsch, G.A.; Rouchka, E.C.; Bhatnagar, A. Identification of a plasma metabolomic signature of thrombotic myocardial infarction that is distinct from non-thrombotic myocardial infarction and stable coronary artery disease. Plos One 2017, 12, e0175591.

7. Jung, J.Y.; Lee, H.S.; Kang, D.G.; Kim, N.S.; Cha, M.H.; Bang, O.S.; Ryu, D.H.; Hwang, G.S. 1h-nmr-based metabolomics study of cerebral infarction. Stroke 2011, 42, 1282–1288.

8. Brereton, R.G.; Lloyd, G.R. Partial least squares discriminant analysis: Taking the magic away. Journal of Chemometrics 2014, 28, 213–225.

9. Gromski, P.S.; Muhamadali, H.; Ellis, D.I.; Xu, Y.; Correa, E.; Turner, M.L.; Goodacre, R. A tutorial review: Metabolomics and partial least squares-discriminant analysis – a marriage of convenience or a shotgun wedding. Analytica Chimica Acta 2015, 879, 10–23.

10. Fahrmann, J.F.; Kim, K.; DeFelice, B.C.; Taylor, S.L.; Gandara, D.R.; Yoneda, K.Y.; Cooke, D.T.; Fiehn, O.; Kelly, K.; Miyamoto, S. Investigation of metabolomic blood biomarkers for detection of adenocarcinoma lung cancer. Cancer Epidemiol Biomarkers Prev 2015, 24, 1716–1723.

11. Frank, I.E.; Friedman, J.H. A statistical view of some chemometrics regression tools. Technometrics 1993, 35, 109.

12. Lê Cao, K.-A.; Martin, P.G.P.; Robert-Granié, C.; Besse, P. Sparse canonical methods for biological data integration: Application to a cross-platform study. BMC Bioinformatics 2009, 10, 34.

13. Lê Cao, K.-A.; Rossouw, D.; Robert-Granié, C.; Besse, P. A sparse pls for variable selection when integrating omics data. Statistical Applications in Genetics and Molecular Biology 2008, 7.

14. Voet, D.; Voet, J.G.; Pratt, C.W. Fundamentals of biochemistry: Life at the molecular level. 4th ed.; Wiley: Hoboken, NJ, 2013.

15. Bishop, C.M. Pattern recognition and machine learning. Springer: New York, 2006; p xx, 738 p.

16. Chih-Wei, H.; Chih-Jen, L. A comparison of methods for multiclass support vector machines. IEEE Transactions on Neural Networks 2002, 13, 415–425.

17. Hammer, B.; Gersmann, K. Neural Processing Letters 2003, 17, 43–53.

18. Hornik, K. Approximation capabilities of multilayer feedforward networks. Neural Networks 1991, 4, 251–257.

19. McCulloch, W.S.; Pitts, W. A logical calculus of the ideas immanent in nervous activity. The Bulletin of Mathematical Biophysics 1943, 5, 115–133.

20. Breiman, L. Random forests. Machine Learning 2001, 45, 5–32.

21. Hastie, T.; Tibshirani, R.; Friedman, J.H. The elements of statistical learning: Data mining, inference, and prediction. 2nd ed.; Springer: New York, NY, 2009; p xxii, 745 p.

22. Breiman, L. Bagging predictors. Machine Learning 1996, 24, 123–140.

23. Tin Kam, H. The random subspace method for constructing decision forests. IEEE Transactions on Pattern Analysis and Machine Intelligence 1998, 20, 832–844.

24. Camacho, D.; de la Fuente, A.; Mendes, P. The origin of correlations in metabolomics data. Metabolomics 2005, 1, 53–63.

25. Steuer, R. Review: On the analysis and interpretation of correlations in metabolomic data. Briefings in Bioinformatics 2006, 7, 151–158.

26. Lewandowski, D.; Kurowicka, D.; Joe, H. Generating random correlation matrices based on vines and extended onion method. Journal of Multivariate Analysis 2009, 100, 1989–2001.

27. Nadarajah, S. A generalized normal distribution. Journal of Applied Statistics 2005, 32, 685–694.

28. Rosipal, R.; Trejo, L.J. Kernel partial least squares regression in reproducing kernel hilbert space. Journal of machine learning research 2001, 2, 97–123.

29. Boulesteix, A.-L. Pls dimension reduction for classification with microarray data. Statistical Applications in Genetics and Molecular Biology 2004, 3, 1–30.

30. Boulesteix, A.L.; Strimmer, K. Partial least squares: A versatile tool for the analysis of high-dimensional genomic data. Briefings in Bioinformatics 2006, 8, 32–44.

31. Höskuldsson, A. Pls regression methods. Journal of Chemometrics 1988, 2, 211–228.

32. Rosipal, R. Nonlinear partial least squares an overview. 2011, 169–189.

33. Lê Cao, K.-A.; Rossouw, D.; Robert-granié, C.; Besse, P. A sparse pls for variable selection when integrating omics data. Statistical Applications in Genetics & Molecular Biology 2008, 7, 1–29.

34. Lê Cao, K.-A.; Boitard, S.; Besse, P. Sparse pls discriminant analysis: Biologically relevant feature selection and graphical displays for multiclass problems. BMC Bioinformatics 2011, 12, 253.

35. Ripley, B.D. Pattern recognition and neural networks. Cambridge University Press: Cambridge ; New York, 1996; p xi, 403 p.

36. Riedmiller, R.; Braun, H. In A direct adaptive method for faster backpropagation learning: The rprop algorithm, IEEE International Conference on Neural Networks, 1993.

37. Vapnik, V.N. Statistical learning theory. Wiley: New York, 1998; p xxiv, 736 p.

38. Thygesen, K.; Alpert, J.S.; Jaffe, A.S.; Simoons, M.L.; Chaitman, B.R.; White, H.D.; Joint, E.S.C.A.A.H.A.W.H.F.T.F.f.U.D.o.M.I.; Authors/Task Force Members, C.; Thygesen, K.; Alpert, J.S., et al. Third universal definition of myocardial infarction. J Am Coll Cardiol 2012, 60, 1581–1598.

